# Reclassification of *Catabacter hongkongensis* as *Christensenella hongkongensis* comb.nov. based on whole genome analysis

**DOI:** 10.1101/2020.09.21.306662

**Authors:** Xiaoying Liu, Jessica L. Sutter, Jacobo de la Cuesta-Zuluaga, Jillian L. Waters, Nicholas D. Youngblut, Ruth E. Ley

**Affiliations:** Department of Microbiome Science, Max Planck Institute for Developmental Biology, Max-Planck-Ring 5, 72076 Tübingen, Germany

**Keywords:** whole genome phylogeny, reclassification, Christensenella, Catabacter

## Abstract

The genera *Catabacter (family Catabacteraceae)* and *Christensenella* (family *Christensenellaceae)* are close relatives within the phylum Firmicutes. Members of these genera are strictly anaerobic, non-spore forming, short straight rods with diverse phenotypes. Phylogenetic analysis of 16S rRNA genes suggest that *Catabacter* splits *Christensenella* into a polyphyletic clade. In an effort to ensure that family/genus names represent monophyletic clades, we performed a whole-genome based analysis of the genomes available for the cultured representatives of these genera: four species of *Christensenella* and two strains of *Catabacter hongkongensis*. A concatenated alignment of 135 shared protein sequences indicates that *C. hongkongensis* is indeed nested within the *Christensenella* clade. Based on their evolutionary relationship, we propose the transfer of *Catabacter hongkongensis* to the new genus as *Christensenella hongkongensis* comb.nov.

## Introduction

*Catabacter hongkongensis* was first isolated in 2007 from the blood cultures of four patients in Hong Kong and Canada. Based on phylogenetic positioning of 16S rRNA sequences and phenotypic characteristics, it was proposed as a new genus and new family, *Catabacteriaceae* [1]. The genus *Catabacter* comprises just one species, with the type strain *Catabacter hongkongensis* HKU16^T^. Based on 16S rRNA gene sequencing surveys, *C. hongkongensis* has been detected in the blood of patients with diseases such as intestinal obstruction, gastrointestinal malignancy, acute cholecystitis and hypertension, in Europe, North America and Asia [1-5].

In 2012, Morotomi and colleagues isolated a novel bacterium from the stool of a healthy male adult, and based on 16S rRNA gene sequence analysis and physiological data, named it *Christensenella minuta* DSM 22607^T^ within the novel family *Christensenellaceae* [6]. In addition to *Christensenella minuta* DSM 22607^T^, three other species have been proposed based on additional isolates from human feces: *Christensenella massiliensis* Marseille-P2438 [7], *Christensenella timonensis* Marseille-P2437 [8], and *Christensenella intestinihominis* AF73-05CM02^PP^ [9]. *Christensenella intestinihominis* AF73-05CM02^PP^ is proposed in a pending patent.

16S rRNA gene sequence identity (%ID) has been used to delineate genus (95 %ID) and species (98.7 %ID) cutoffs [10, 11]. The 16S rRNA gene sequence of *C. hongkongensis* HKU16^T^ has 96-97 %ID with the 16S rRNA genes of the four species of *Christensenella*, which places them in the range of sharing a genus using that criterion. In addition to sequence similarity, the 16S rRNA gene-based phylogenetic relationships of these taxa indicate they form a monophyletic clade [12].

Whole genome-based analysis with concatenated protein sequences has recently replaced 16S rRNA-based phylogenetics as a basis for determining the evolutionary history of members of the Bacteria and Archaea [13]. Based on whole genome comparisons, *Catabacter* and *Christensenella* were annotated as belonging to the family *Christensenellaceae* in the order *Christensenellales* in the Genome Taxonomy Database (GTDB; R05-RS95 17^th^ July 2020) [14]. Twenty-one genomes within the family *Christensenellaceae* are included in GTDB R05-RS95 as of 01 August 2020. These include metagenome-assembled genomes and genomes derived from isolates. A formal reclassification of *Catabacter* as *Christensenella* would clarify the nomenclature of this taxon.

Here, we used comparative genomics as a basis for proposing that the genus name *Catabacter* and the family name Catabacteraceae be removed from the nomenclature. Genome sequences of six cultured isolates belonging to the families *Catabacteriaceae* and *Christensenellaceae* and four species from sister clades in GTDB were selected for phylogenomic analysis. The average nucleotide identity (ANI) of the six genomes in the family of *Catabacteriaceae* and *Christensenellaceae* were compared. Based on the resulting phylogeny, we recommend that *Catabacter hongkongensis* be renamed *Christensenella hongkongensis* comb.nov.

## Methods

### Phylogeny based on whole genomes

We based this analysis on whole genome sequences of six cultured isolates: *Catabacter hongkongensis* strains HKU16^T^ and ABBA15k, *Christensenella minuta* DSM 22607^T^, *Christensenella massiliensis* Marseille-P2438, *Christensenella timonensis* Marseille-P2437, *Christensenella intestinihominis* AF73-05CM02^PP^. The general information about genomes in this study is listed in Table 1. In addition, we selected for the outgroup the species *Clostridium novyi* NT (GenBank accession number: GCA_000014125.1), *Clostridium butyricum* DSM 10702^T^ (GenBank accession number: GCA_000409755.1), *Clostridium thermobutyricum* DSM4928^T^ (GenBank accession number: GCA_002050515.1) and *Eubacterium limosum* ATCC 8486^T^ (GenBank accession number: GCA_000807675.2). Whole genome sequences were obtained from NCBI.

**Table 1.** Phenotypic characteristics of the strains of *Catabacter* and *Christensenella* based on literature review. Data for the strains are from references [1, 6-9, 30]. +, Positive; -, negative; ND, not determined. The G+C contents and N50, contig numbers, genome size and genome coverages were retrieved from the GTDB records of the strains.

| Characteristics | *Christensenella minuta*<br>DSM 22607<sup>T</sup> | *Christensenella intestinhominis*<br>AF73-05CM02<sup>PP</sup> | *Christensenella massiliensis*<br>Marseille-P2438 | *Christensenella timonensis*<br>Marseille-P2437 | *Catabacter hongkongensis* |  |
| --- | --- | --- | --- | --- | --- | --- |
|  |  |  |  |  | HKU16<sup>T</sup> | ABBA15k |
| Gram stain | Negative/Positive | Positive | Negative | Negative | Positive | Positive |
| Motility | Nonmotile | Nonmotile | Nonmotile | Nonmotile | Motile | Nonmotile |
| Catalase activity | - | - | - | - | + | ND |
| Metabolite utilization | Arabinose, Glucose, Mannose, Rhamnose, Salicin, Xylose | Arabinose, Glucose, Mannose, Rhamnose, Xylose, Mannitol, Maltose, Sulphate, Pine syrup, Raffinose, Sorbitol | ND | ND | Arabinose, Glucose, Mannose, Xylose | ND |
| G+C content (%) | 51.48% | 52.07% | 50.38% | 51.71% | 48.53% | 48.79% |
| Contig number | 45 | 36 | 1 | 2 | 134 | 113 |
| Protein count | 2776 | 2791 | 2437 | 2430 | 3071 | 2625 |
| Completeness (Contamination) | 98.39% (0.81%) | 99.19% (0.81%) | 98.79 % (0.81%) | 97.98% (0.81%) | 97.55% (2.97%) | 97.9% (3.5%) |
| Genome size | 2,940,227 bp | 3,026,655 bp | 2,560,186 bp | 2,650,850 bp | 3,203,641 bp | 2,797,114 bp |
| GenBank Assembly Accession | GCA_001678855.1 | GCA_001678845.1 | GCA_900155415.1 | GCA_900087015.1 | GCA_000981035.1 | GCA_001507385.1 |

We used Anvi’o v5.2.0 for constructing the whole-genome phylogenomic tree [15]. Briefly, contig databases were created from the genome FASTA files. Prodigal v2.6.3 with default settings [16] was used to identify open reading frames in contigs. Hidden Markov model (HMM) profiles were used to extract the set of single-copy marker genes defined by Campbell et al. [17]. The best HMM hit was selected if a gene was found with multiple copies in a genome. We limited the set of single-copy core genes shared to those present in all analyzed genomes and aligned the concatenated protein sequences using muscle [18].

FastTree 2 [19] was used for constructing approximately-maximum-likelihood phylogenomic tree with the Jones-Taylor-Thornton model [20]. SH-like local support values [21] are shown on the nodes. The phylogenetic tree was visualized by using the online tool iTOL [22].

### Average nucleotide identity and phenotype predictions

We used FastANI with default settings [23] to generate a pairwise ANI comparison of the six *Christensenella* and *Catabacter* genomes. A heatmap of ANI values was generated and visualized in R [24] with the package ggplot2 [25]. Traitar [26] trait analyzer was used for phenotypic trait prediction based on genome sequences. ABRicate v1.0.1 (https://github.com/tseemann/ABRicate) was used for the detection of genes involved in antimicrobial resistance (AMR), and the annotation was derived from the default NCBI database AMRFinderPlus.

## Results

The genome sizes of the six *Catabacter* and *Christensenella* species/strains range from 2.5 Mbp to 3.3 Mbp and the G+C content of genomic DNA from 48.53 to 52.07 %. Based on the pairwise comparison of the six genomes in the family of *Catabacteriaceae* and *Christensenellaceae*, we observed that the ANI of the two *Catabacter hongkongensis* strains (HKU16^T^ and ABBA15k) was >98.97 % (Fig. 1), confirming that the two strains belong to the same species. Moreover, the ANI values for the six genomes were between 77.56-83.48 %, which corresponds to the accepted ANI cut-off 94-96 % used to designate the same species [27-29] and <83 % for inter-species ANI values [23]. *Christensenella intestinihominis* AF73-05CM02^PP^ and *C. minuta* DSM 22607^T^ showed the highest ANI similarity values (83.48%) between different species.

**Fig. 1.**
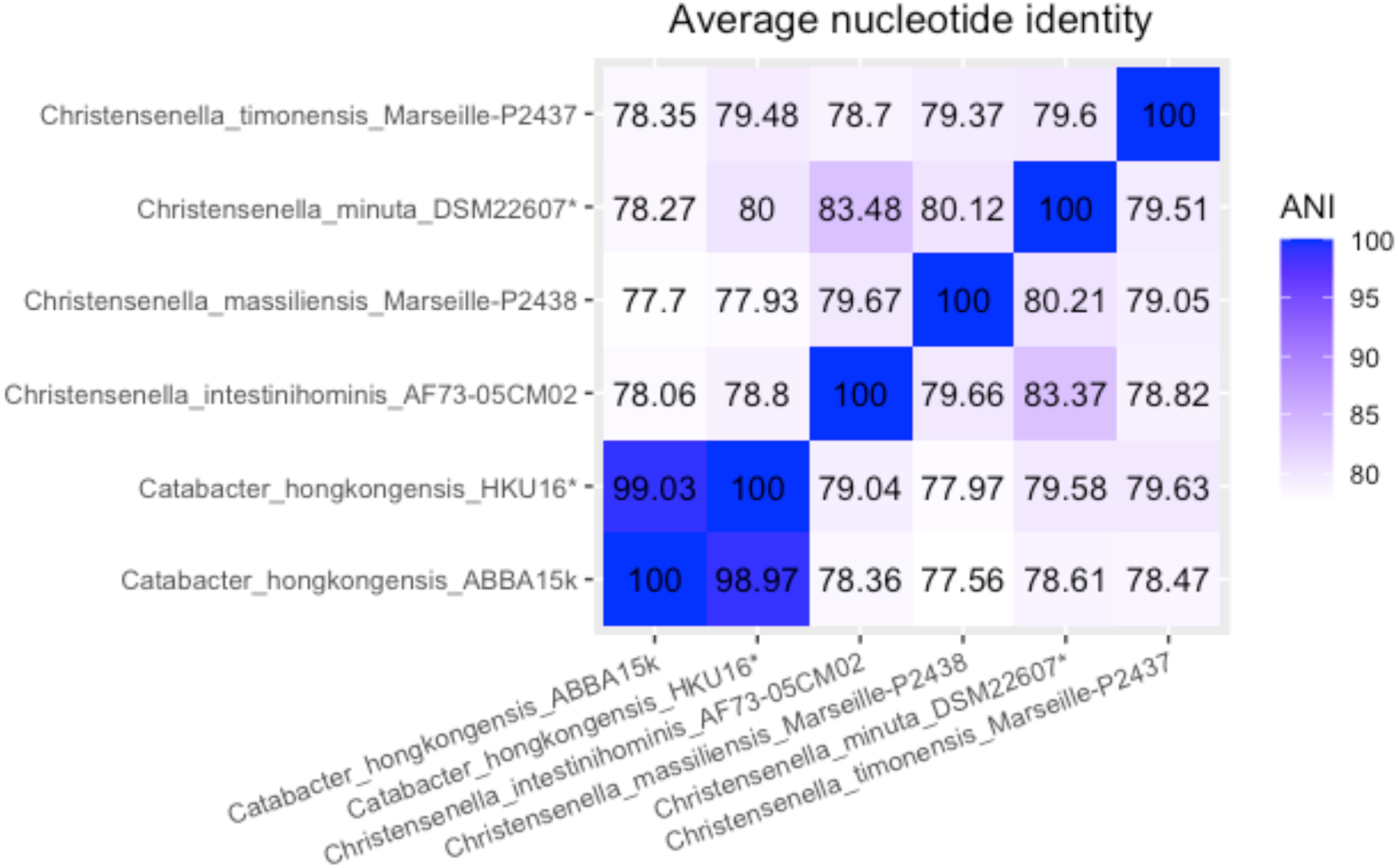
Heatmap of ANI values amongst the genomes of *Catabacter hongkongensis* strains and *Christensenella* species in this study. Type strains are marked with asterisk.

We identified 135 protein-encoding single-copy core genes present in the genomes of *Christensenella, Catabacter* and the outgroup taxa. We used these 135 genes in a concatenated alignment resulting in a total of 51,813 aligned amino acid sites. In the resulting phylogenetic tree (Fig. 2), the *Catabacter* and *Christensenella* species and strains formed a monophyletic clade with high bootstrap support, indicating a shared common ancestor. The species *C. timonensis* Marseille-P2437 is basal and forms a sister clade to the rest of the taxa in the phylogeny. The two strains of *Catabacter hongkongensis* (HKU16^T^ and ABBA15k) are, as expected based on their high ANI, on the same branch of the phylogeny. The *Catabacter* branch is a sister taxon to the remaining *Christensenella* species (*C. minuta* DSM 22607^T^, *C. massiliensis* Marseille-P2438, *C. intestinihominis* AF73-05CM02^PP^).

**Fig. 2.**
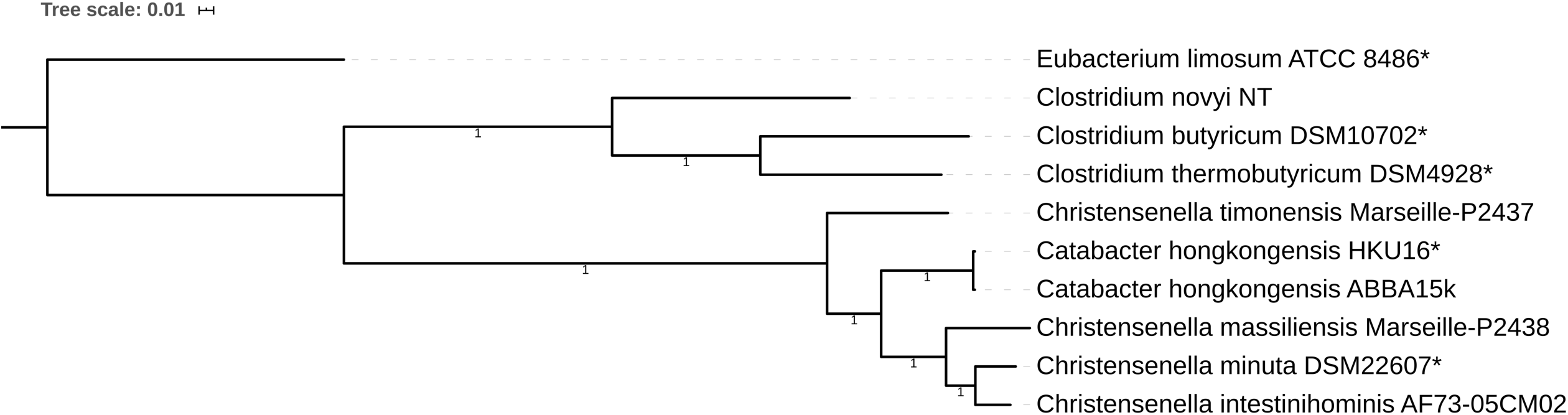
Phylogeny reconstructed by approximately-maximum-likelihood showing the position of *Catabacter* relative to *Christensenella* based on 135 concatenated core protein sequences with 51,813 aligned amino acid sites. All the nodes are strongly supported with SH-like support values of 1. Type strains are marked with asterisk. *Clostridium* and *Eubacterium* are used for the outgroup. The tree is rooted by *Eubacterium limosum* ATCC 8486. Scale bar indicates 0.01 amino acid substitutions per site.

The position of *Catabacter* (and its family Catabacteriaceae) nested within the *Christensenella* clade splits the Christensenellaceae family and genus, such that neither are monophyletic. For the family and genus names to represent monophyletic groups the renaming of *Catabacter hongkongensis* to *Christensenella hongkongensis* would be required. As a consequence, the genus name Catabacter and Catabacteriaceae should be removed from the nomenclature.

The cultured strains of the species of *Catabacter* (*C. hongkongensis* HKU16^T^ and ABBA15k) and *Christensenella* (*C. minuta* DSM 22607^T^, *C. massiliensis* Marseille-P2438, *C. timonensis* Marseille-P2437, *C. intestinihominis* AF73-05CM02^PP^) have been shown to be strictly anaerobic and non-spore forming rods with varied motility, Gram stain reaction and the catalase reaction [1, 6-9]. The different phenotypic characteristics of the species compared in this study is summarized in Table 1. *Catabacter hongkongensis* HKU16^T^ and ABBA15k strains are reported to be Gram-positive, while the four species of *Christensenella* are reported either Gram-positive or Gram-negative. Morotomi and colleagues reported that C. *minuta* DSM 22607^T^ is Gram-negative [6], while another group reports *C. minuta* stains consistently as Gram-positive [30], which is consistent with our observations [12]. The Gram-variable reaction might be due to the age of the culture for staining [31]. However, based on the phenotype predictions of the included genomes by using Traitar trait analyzer, all of the strains in those two genera are predicted to produce a cell wall that would be consistent with a Gram-positive reaction.

*C. hongkongensis* strains (HKU16^T^, HKU17, CA1, CA2) and most clinical derived isolates are reported to be motile and resistant to cefotaxime [1, 2, 5, 32] except for *C. hongkongensis* ABBA15k, which was isolated in 2016 from the blood of a patient with a fever in Sweden [33]. Strain ABBA15k showed 100% 16S rRNA gene identity with *Catabacter hongkongensis* HKU16^T^. However, the genome of *C. hongkongensis* ABBA15k is smaller than *C. hongkongensis* HKU16^T^, and the genes coding for chemotaxin (*cheA*) and flagellar assembly (*flhA* and MotA) were not present in the genome of *C. hongkongensis* ABBA15k [33]. The tetracycline resistance gene *tet* was detected in the genome of *C. hongkongensis* HKU16T, but no resistance genes were detected in the genome of *C. hongkongensis* ABBA15k [33].

Screening for AMR genes of the genomes with ABRicate in this study showed that the *tet* gene was also present in the genomes of *Christensenella minuta* DSM 22607^T^, *Christensenella massiliensis* Marseille-P2438, *Christensenella timonensis* Marseille-P2437 and *Catabacter hongkongensis* HKU16T but not in *Christensenella intestinihominis* AF73-05CM02^PP^ and *Catabacter hongkongensis* ABBA15k. A Streptomycin resistance gene (*aadE*) was also detected in the genome of *Christensenella massiliensis* Marseille-P2438. The detailed information about AMR genes is listed in Table 2. *Christensenella intestinihominis* AF73-05CM02^PP^ and *Catabacter hongkongensis* HKU16^T^ were predicted to be motile by Traitar. However, *Christensenella intestinihominis* AF73-05CM02^PP^ was classified as non-motile in the original phenotypic characterization [9], which might be attributable to the growth conditions used. It is also possible that the genome of the strain may not contain all genes required for flagellar formation.

**Table 2.** Antimicrobial resistance genes (AMR) detected for the genomes of *Catabacter hongkongensis* strains and *Christensenella* species. Coverage refers to the proportion of the gene in the reference gene sequence.

| Strain | Contig<br>(Position Strand) | Reference Gene<br>(Accession) | Coverage | Identity % | Gene product | Resistance |
| --- | --- | --- | --- | --- | --- | --- |
| *Christensenella timonensis*<br>Marseille-P2437 | FLKP01000002.1<br>(1477797–1479716 +) | *tet(W)*<br>(NG_048299.1) | 1-1920/1920 | 99.53 | tetracycline resistance<br>ribosomal protection<br>protein Tet(W) | TETRACYCLINE |
|  | FLKP01000002.1<br>(1480702–1481922 +) | *tet(40)*<br>(NG_048141.1) | 1-1221/1221 | 99.67 | tetracycline efflux MFS<br>transporter Tet(40) | TETRACYCLINE |
| *Christensenella massiliensis*<br>Marseille-P2438 |  | *tet(W)*<br>(NG_048281.1) | 1-1920/1920 | 100 | tetracycline resistance<br>ribosomal protection<br>protein Tet(W) | TETRACYCLINE |
|  | LT700187.1<br>(1980989–1981855 +) | *aadE*<br>(NG_047378.1) | 1-867/867 | 99.77 | aminoglycoside 6-adenylyltransferase<br>AadE | STREPTOMYCIN |
| *Catabacter hongkongensis*<br>HKU16<sup>T</sup> | LAYJ01000061.1<br>(37275–39194 +) | *tet(32)*<br>(NG_048125.1) | 1-1920/1920 | 100 | tetracycline resistance<br>ribosomal protection<br>protein Tet(32) | TETRACYCLINE |
| *Christensenella minuta*<br>DSM 22607<sup>T</sup> | MAIR01000011.1<br>(54376-56295 +) | *tet(W)*<br>(NG_048281.1) | 1-1920/1920 | 100 | tetracycline resistance<br>ribosomal protection<br>protein Tet(W) | TETRACYCLINE |

In conclusion, both *Catabacter* and *Christensenella* genera include species and strains that are strictly anaerobic, non-spore forming, short straight rods and have diverse phenotypes regarding motility, Gram-staining and antibiotic resistance. The genus *Catabacter* was proposed earlier, however, only one species exists within in genus and the family Catabacteriaceae, while four species have been proposed for the genus *Christensenella* and the family Christensenellaceae. Based on previously reported pairwise 16S rRNA gene sequence identities and our genome-based phylogenomic analysis, we propose that the genus *Catabacter* and the family Catabactericaeae be removed from the nomenclature and that the species *Catabacter hongkongensis* be renamed *Christensenella hongkongensis* comb.nov.

### Description of *Christensenella hongkongensis* comb.nov

The description of *Christensenella hongkongensis* is identical to that proposed for *Catabacter hongkongensis* [1].

The type strain is HKU16^T^ (= DSM 18959^T^ = JCM 17853^T^ = CCUG 54229^T^).

## Acknowledgements

This research was supported by the Max Planck Society.

